# Protective porcine influenza virus-specific monoclonal antibodies recognize similar haemagglutinin epitopes as humans

**DOI:** 10.1101/2020.07.21.213470

**Authors:** Barbara Holzer, Pramila Rijal, Adam McNee, Basu Paudyal, Becky Clark, Tanuja Manjegowda, Francisco J. Salguero, Emily Bessell, John C. Schwartz, Katy Moffat, Miriam Pedrera, Simon P. Graham, Marie Bonnet-Di Placido, Roberto M. La Ragione, William Mwangi, Peter Beverley, John W. McCauley, Rodney S. Daniels, John Hammond, Alain R. Townsend, Elma Tchilian

## Abstract

Pigs are natural hosts for the same subtypes of influenza A viruses as humans and integrally involved in virus evolution with frequent interspecies transmissions in both directions. The emergence of the 2009 pandemic H1N1 virus illustrates the importance of pigs in evolution of zoonotic strains. Here we generated pig influenza-specific monoclonal antibodies (mAbs) from H1N1pdm09 infected pigs. The mAbs recognized the same two major immunodominant haemagglutinin (HA) epitopes targeted by humans, one of which is not recognized by post-infection ferret antisera that are commonly used to monitor virus evolution. Neutralizing activity of the pig mAbs was comparable to that of potent human anti-HA mAbs. Further, prophylactic administration of a selected porcine mAb to pigs abolished lung viral load and greatly reduced lung pathology but did not eliminate nasal shedding of virus after H1N1pdm09 challenge. Hence mAbs from pigs, which target HA can significantly reduce disease severity. These results, together with the comparable sizes of pigs and humans, indicate that the pig is a valuable model for understanding how best to apply mAbs as therapy in humans and for monitoring antigenic drift of influenza viruses in humans, thereby providing information highly relevant to making influenza vaccine recommendations.

---

Pigs are natural hosts for influenza A viruses (IAV) and closely related H1N1 and H3N2 viruses circulate in pigs and humans^1^. Frequent interspecies transmissions between pigs and humans contributes to the evolution of IAV and can be a source for novel pandemic strains^2-4^. Pigs are anatomically, physiologically and immunologically more similar to humans than laboratory animals (mice, guinea pigs and ferrets) commonly used for influenza virus research^5,6^. Pigs and humans have similar distributions of sialic acid receptors in their respiratory tracts and longer life spans than laboratory animals which, with their more comparable size, makes them a useful stepping stone for translation of experimental results into human clinical applications. Furthermore, the dynamics of IAV transmission in pigs are well characterized and many immunologic tools, such as inbred strains, tetramers and antibodies for surface markers and cytokines are available to dissect porcine immune responses^7^.

In recent years monoclonal antibodies (mAbs) from mice and humans have been used to investigate the antigenic structure of the influenza haemagglutinin (HA) glycoprotein and mechanisms of escape from immune surveillance. Three main classes of anti-HA antibodies have been defined. The majority of most potent neutralizing mAbs are targeted to the HA head and they are subtype and clade specific^8,9^. A second class of HA head mAbs are broadly neutralizing within the subtype, while the third category is broadly neutralizing across HA subtypes and they mainly target the HA stem^10-14^. In the last decade the anti-stem mAbs have received considerable attention as a possible therapy for severe influenza disease, but so far success in small animal models has not been translated into success in the clinic^15-18^. We have shown that a strongly neutralizing human mAb, 2-12C, against the HA head, administered prophylactically to pigs reduced virus shedding and lung pathology after influenza challenge. Similarly, administration of DNA encoded 2-12C mAb has shown promise as a novel therapeutic strategy by reducing lung pathology in pigs^19^. The therapeutic administration of the human broadly neutralizing anti-stem mAb, FI6, did not reduce virus load in pigs, although it did reduce lung pathology^20^.

However, longer term investigation of human mAbs in pigs is compromised by development of anti-human Ig responses. To overcome this limitation, we developed porcine mAbs to pandemic H1N1 influenza virus which will increase the utility of the pig model in influenza virus research for evaluation of therapeutic mAbs and delivery platforms

## Results

### Generation of porcine H1N1pdm09-specific mAbs

To generate a high frequency of antibody secreting cells, two pigs were challenged intranasally with a swine H1N1 isolate, derived from the A(H1N1)pdm09 virus (H1N1pdm09) and re-challenged 22 days post infection (DPI) with the same virus. Both pigs shed virus after the first challenge, confirming successful influenza infection, but shedding was not detected after the second challenge (**Fig. 1a**). Serum neutralizing antibody titres to H1N1pdm09 and binding to HA from A/England/195/2009 (pHA) were determined at 7, 14, 22 and 29 DPI, showing that both pigs developed strong antibody responses with neutralizing titer of 1:640 (**Figs. 1b and c**). Analysis by *ex vivo* B cell ELISpot revealed that the number of HA-specific IgG-secreting cells in the blood was low compared to tracheobronchial lymph nodes (TBLN), while lung, tonsil and broncho-alveolar lavage (BAL) tissues had intermediate levels (**Fig. 1d**). Therefore, we used TBLN from pig 12 at day 7 post re-challenge, to sort HA-specific B cells using biotinylated pHA. The gating strategy is shown in **Supp Fig 1**.

**Figure 1.**
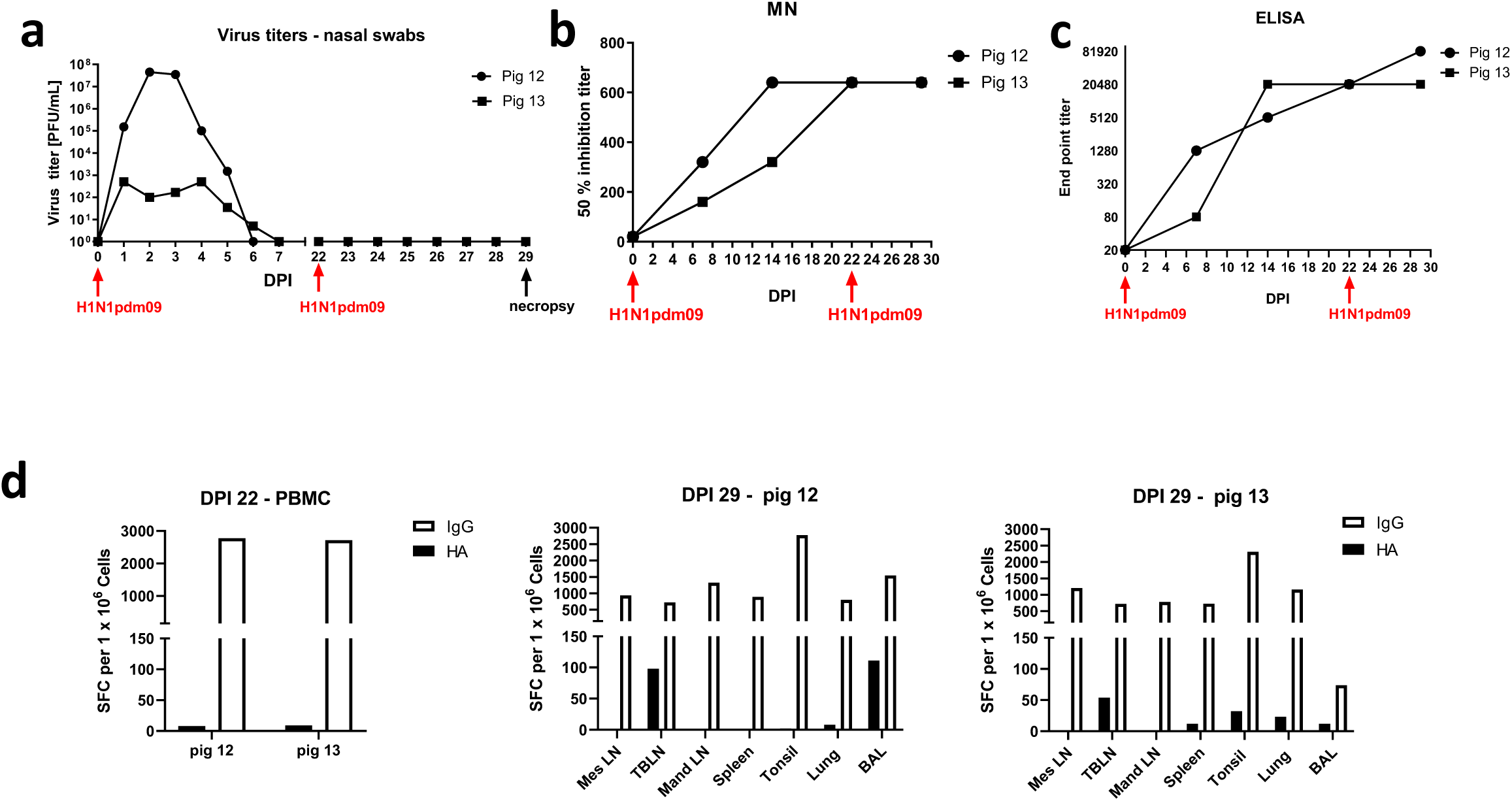
Responses in H1N1pdm09 infected pigs used for mAb generation. Two pigs (pig12 and pig13) were challenged intranasally with H1N1pdm09 and 22 days later re-challenged with the same virus. Virus loads in nasal swabs, taken daily after challenge, were assessed by plaque assay (**a**), serum neutralizing activity of H1N1pdm09 by microneutralization assay (**b**) and pHA binding was quantified by ELISA (**c**). pHA specific and porcine IgG producing spot forming cells (SFC) were enumerated in blood at 22 DPI and in mesenteric (Mes LN), tracheobronchial (TBLN), mandibular (Mand LN), spleen, tonsils, lung and broncho-alveolar lavage (BAL) at 29 DPI (7 days post re-challenge) (**d**).

A total of 70 single cells were sorted and 45 were positive for the heavy and light chains of IgG (63% recovery). cDNA was synthesized followed by a nested PCR amplification to generate gamma and kappa chains for cloning, and a single PCR step for the production and cloning of the lambda chains. PCR products were cloned into porcine expression cassettes containing constant domains of the kappa, lambda and gamma chains. All amplified gamma heavy chains were IgG1.

Of the 45 antibody pairs 19 had a lambda light chain and 26 had kappa light chains (**Table 1**). The germline light chain V gene segments are well annotated for the pig but are polymorphic. Therefore it is difficult to distinguish between closely related gene segments due to V(D)J recombination yielding imprecise junctions and the presence of somatically hypermutated clones^21-24^. Presumably because of these processes, none of the light chains were identical and were solely classified by subfamily (e.g. IGKV1, IGKV2, IGLV3, and IGLV8). The porcine IGH region is incompletely annotated, polymorphic, and the genes tend to be highly similar (same subfamily) outside the CDRs and often share CDRs between genes and alleles ^25,26^. It is therefore difficult to determine IGHV gene segment usage (**Table 1**).

**Table 1.** Characteristics of porcine anti-influenza mAbs.

| Clone | ELISA<br>pHA | MN pH1N1<br>(ng/ml) | HAI<br>(ng/ml) | Light chain V<br>family | CDRH3 length<br>(aa) | Cysteines in<br>CDRH3 |
| --- | --- | --- | --- | --- | --- | --- |
| pb1 | Yes | 5 | 312.5 | IGKV2 | 12 |  |
| pb2 | Yes | No | No | IGLV8 | 19 |  |
| pb3 | Yes | 20,480 | 50,000 | IGKV2 | 17 |  |
| pb4 | Yes | No | No | IGLV8 | 12 |  |
| pb5 | Yes | No | No | IGKV1 | 15 |  |
| pb6 | No | No | No | IGLV3 | 16 | 2 |
| pb7 | No | No | No | IGLV3 | 19 | 2 |
| pb8 | Yes | No | No | GKV2 | 16 |  |
| pb9 | Yes | No | No | IGLV8 | 20 | 2 |
| pb10 | Yes | 5,120 | 12,500 | IGLV8 | 22 | 2 |
| pb11 | Yes | 80 | 625 | IGLV8 | 10 |  |
| pb12 | Yes | No | No | IGLV8 | 20 | 2 |
| pb13 | Yes | 10,240 | 25,000 | IGLV8 | 8 |  |
| pb14 | Yes | 5 | 156.25 | IGLV8 | 21 | 2 |
| pb15 | Yes | 20 | 312.5 | IGLV8 | 21 |  |
| pb16 | Yes | 20 | 312.5 | IGLV8 | 17 | 2 |
| pb17 | Yes | No | No | IGLV8 | 20 | 2 |
| pb18 | Yes | 10 | 156.25 | IGLV8 | 18 | 2 |
| pb19 | Yes | 5,120 | 25,000 | IGLV8 | 24 | 2 |
| pb20 | Yes | 10 | 156.25 | IGLV8 | 18 | 2 |
| pb21 | Yes | No | No | IGLV8 | 19 |  |
| pb22 | Yes | No | No | IGKV1 | 8 |  |
| pb23 | Yes | No | No | IGKV1 | 14 |  |
| pb24 | Yes | 20 | 625 | IGKV2 | 12 |  |
| pb25 | Yes | No | No | IGKV1 | 13 |  |
| pb26 | No | No | No | IGKV2 | 19 |  |
| pb27 | Yes | 10 | 156.25 | IGKV2 | 13 |  |
| pb28 | Yes | No | No | IGKV2 | 19 |  |
| pb29 | Yes | 40,960 | 100,000 | IGKV1 | 20 |  |
| pb31 | Yes | No | No | IGKV1 | 17 |  |
| pb32 | No | No | No | IGKV2 | 12 |  |
| pb33 | Yes | No | No | IGKV2 | 15 |  |
| pb34 | Yes | No | No | IGKV1 | 15 |  |
| pb35 | Yes | No | No | IGKV2 | 20 |  |
| pb36 | Yes | No | No | IGKV2 | 15 |  |
| pb37 | Yes | No | No | IGKV1 | 12 |  |
| pb38 | Yes | No | No | IGKV2 | 13 |  |
| pb39 | Yes | 5,120 | 12,500 | IGKV2 | 18 |  |
| pb40 | Yes | No | No | IGKV2 | 14 |  |
| pb41 | Yes | No | No | IGKV1 | 15 | 2 |
| pb42 | No | No | No | IGKV1 | 15 |  |
| pb43 | Yes | 10,240 | 50,000 | IGKV2 | 13 |  |
| pb44 | Yes | No | No | IGLV8 | 8 |  |
| pb45 | Yes | 5,120 | 6,250 | IGLV8 | 21 | 2 |
| pb46 | Yes | 10 | 312.5 | IGKV2 | 19 |  |

**Table 2.** Amino acid substitutions selected by antibodies in the HA of H1N1pdm09 virus. Mutant viruses selected with pb1, pb20 and pb24 encoded HA homogenous amino acid substitutions. Those selected with pb15, pb18 and pb27 encoded polymorphisms at two or more amino acid positions; ratios of parent/mutant amino acids at each position are indicated.

| Mab | HA amino acid substitutions |  |  |  |  |  |  |
| --- | --- | --- | --- | --- | --- | --- | --- |
|  | K130 | G155 | K160 | G170 | A186 | D187 | Q223 |
| **pb1** |  | E |  |  |  | E | R |
| **pb15** | K20/E80 |  |  | G80/R20 |  |  |  |
| **pb18** |  |  | K45/E55 |  | A35/D65 |  |  |
| **pb20** |  |  |  |  | D |  |  |
| **pb24** |  | E |  |  |  | E | R |
| **pb27** | K45/E55 | G70/E30 |  |  |  | D70/E30 | Q70/R30 |

Each native pair of light and heavy chain expressing plasmids was transiently transfected into HEK293 cells to produce recombinant porcine mAbs and the IgGs were purified from culture-supernatants. All 45 mAbs expressed and 40 bound to pHA.

### Neutralizing and binding activity of porcine influenza mAbs

Of the 40 mAbs binding pHA in ELISA, nine strongly neutralized H1N1pdm09. The mean concentration giving 50% inhibition was 12.2 ng/ml (range 2.5 to 20 ng/ml) which is comparable to the strongly neutralizing human mAb 2-12C (20 ng/ml)^19,27^. Among other mAbs pb11 inhibited at 80 ng/ml and clones pb3, pb10, pb13, pb19, pb39, pb43 and pb45 at greater than 5 μg/ml (**Fig. 2a and Table 1**). mAb pb29 inhibited at ∼41 μg/ml. The mAbs had a similar pattern of hemagglutination inhibition (HAI) to their neutralization (**Fig. 2b and Table 1**).

**Figure 2.**
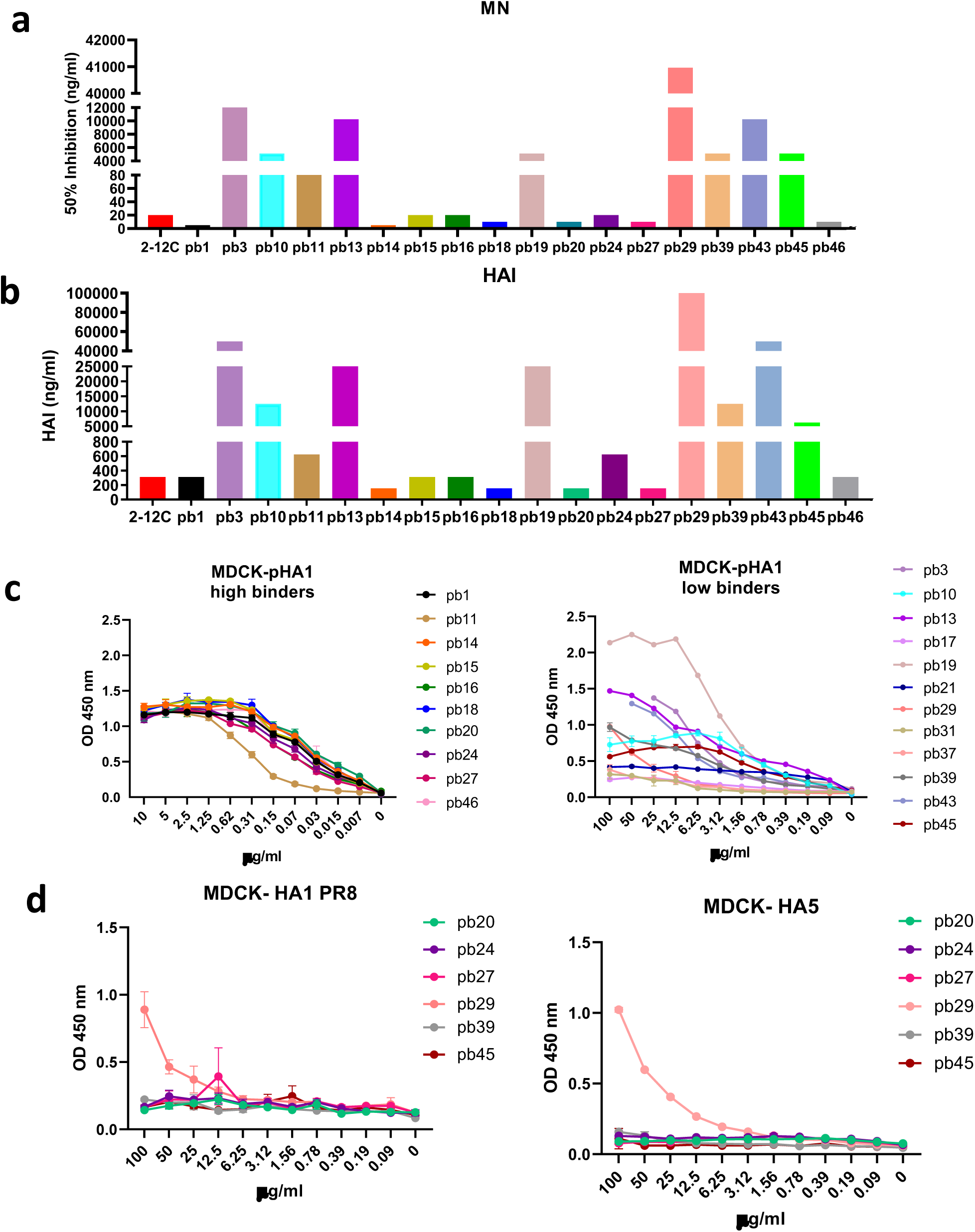
Neutralizing and binding activity of porcine mAbs. Concentrations of individual mAbs giving 50% neutralization (**a**) and hemagglutination inhibition (HAI) (**b**). Titration of binding of porcine mAbs to MDCK-pHA (**c**) and MDCK HA PR8 and MDCK-HA 5 expressing cells (**d**).

We also tested the binding of the mAbs to MDCK-SIAT1 cells stably expressing pHA. The binding strength corelated positively with the neutralization potency. The 9 strongly neutralizing mAbs show saturation binding in the range of 0.3 to 0.6 μg/ml, while pb11 saturated at a slightly higher concentration of 1.25 μg/ml. The poorly neutralizing pb3, pb10, pb13, pb19, pb29, pb39, pb43 and pb45 saturate at higher concentrations of 6.2 to 12.5 μg/ml (**Fig. 2c**). The rest of the non-neutralizing mAbs showed weak or no binding to the cells at very high concentrations despite all of them being positive in ELISA. This may be due to the native HA on the cell surface as opposed to the secreted protein. Antibody pb29 also bound to MDCK cells expressing HAs from A/Vietnam/1203/2004 (A(H5N1)) and PR8 (A(H1N1)) viruses (**Fig. 2d**). There was no correlation between IGHV, IGKV, or IGLV gene usage or CDR-H3 length with neutralization or binding.

We analyzed the inhibition of binding of the human anti-head mAbs 2-12C and T3-4B, and the anti-stem mAb MEDI8852 to MDCK-SIAT1-pHA by the porcine mAbs ^27^. Only the porcine mAbs pb1, pb14, pb15, pb18, pb20, pb24 and pb27 blocked binding of the anti-head mAbs and these pig mAbs are therefore considered to bind to HA head (**Fig. 3a**). These antibodies also inhibited hemagglutination (**Fig. 3b**). However, it is difficult to allocate them to a specific epitope within the head because of the steric hindrance by IgG. In contrast, pb29 blocked binding of MEDI8852, did not inhibit hemagglutination and was cross-reactive with HAs of other H1 and H5 subtypes (**Fig. 2d**), indicating that this might be a stem-specific mAb.

**Figure 3.**
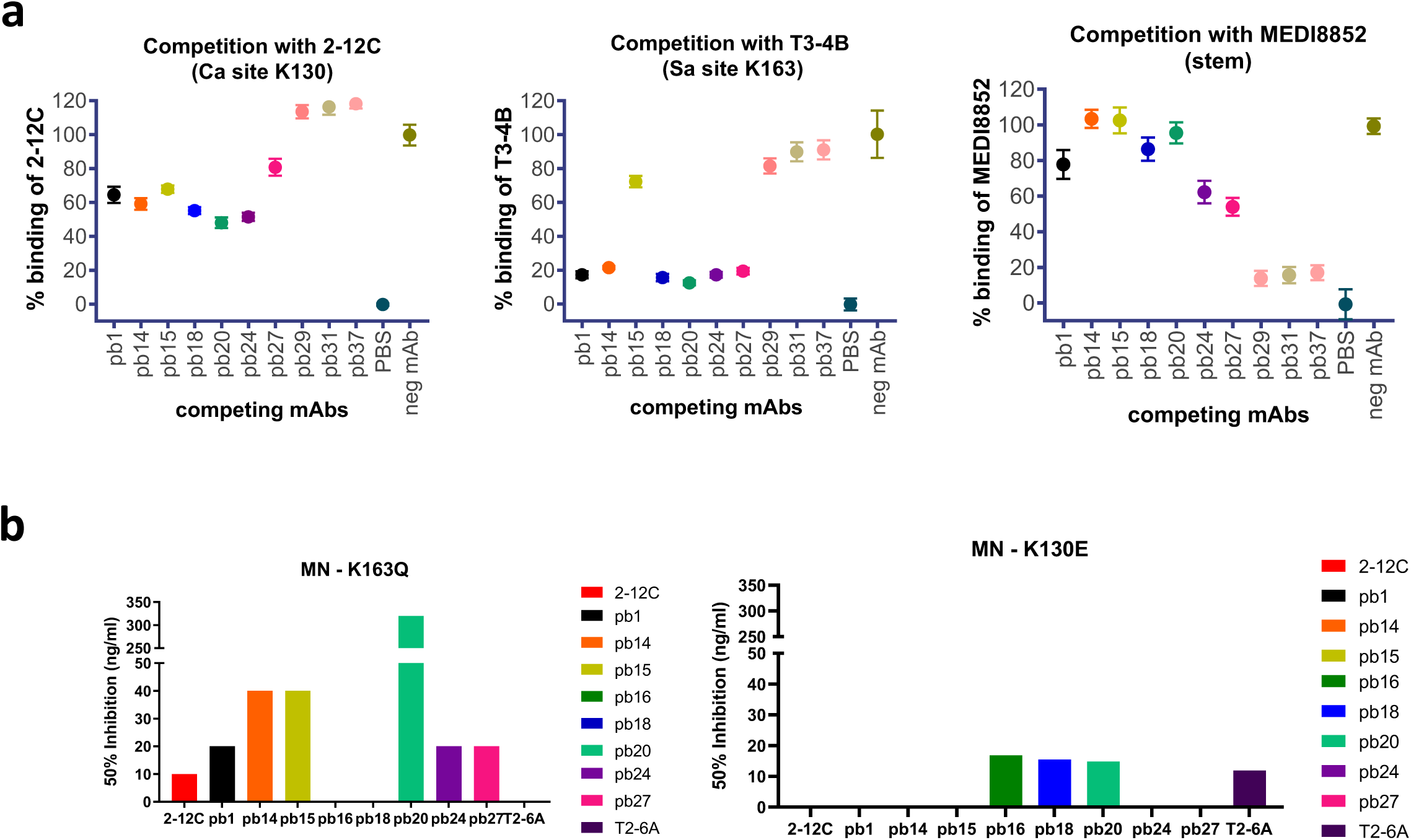
Inhibition of binding of human mAbs and neutralization of mutant viruses by porcine mAbs. Inhibition of binding of human 2-12C, T3-4B and MEDI885 mAbs to MDCK-pHA expressing cells by porcine mAbs. Means and 95% confidence intervals for 8 replicates are shown (**a**). Percent competition was calculated as (X-Max)/(Max-Min)*100, where X= Measurement value, Max = Negative mAb (EBOV Mab), Min = PBS. Concentrations of mAbs giving 50% neutralization of viruses with HA K163Q and K130E substitutions (**b**). Absence of bar indicates that a mAb is non-neutralizing.

### HA epitope recognition

We tried to define the HA sites recognised by the mAbs by analyzing the neutralization of a H1N1pdm09 vaccine virus X-179A (with HA and NA of A/California/07/2009 virus) containing a K163Q HA1 amino acid substitution. This lies in the Sa antigenic site of the pHA and is the defining substitution of antigenically drifted clade 6B human influenza viruses that spread globally in 2014^27^. Antibodies pb1, pb14, pb15, pb24, pb27 and the human 2-12C all neutralized K163Q variant virus with 50% inhibition at concentrations between 10 and 40 ng/ml. pb20 required a higher concentration of 320 ng/ml for neutralization (**Fig. 3b**). mAbs pb16, pb18 and the control human mAb T2-6A did not neutralize virus with the K163Q substitution. However, pb16, pb18 and pb20 neutralized virus with a HA1 K130E substitution at 10 to 15 ng/ml, while pb1, pb14, pb15, pb24 and 2-12C did not.

To identify the site recognised by mAbs pb1, pb15, pb18, pb20, pb24 and pb27 we selected escape mutants of the X-179A virus. Gene sequencing was performed on escape mutant populations, not single clones, hence substitutions in multiple sites in the HA head were observed but no accompanying substitutions in the NA were found. Antibodies pb1, pb15, pb24 and pb27 are classified to be of one group where they selected one or more substitutions in the Ca site (K130E, Q223R), the Sa site (G155E) and the Sb site (D187E) (**Fig. 4**). These substitutions surround the sialic acid binding site but not all are necessarily key substitutions that alter virus antigenicity; G155E and Q223R are often selected as cell culture- or egg-propagation adaptations. The escape mutants were either not neutralized or required higher concentrations of mAbs to neutralize. The key antigenic substitution, K130E, was selected by pb15 and pb27, as it was by the human mAb 2-12C ^27^. Two other mAbs, pb18 and pb20, selected substitutions in the Sa site (K160E) and the Sb site (A186D) and did not neutralize the K163Q X-179A virus, making them similar to the previously described human Sa site mAbs T3-4B and T2-6A^27^.

**Figure 4.**
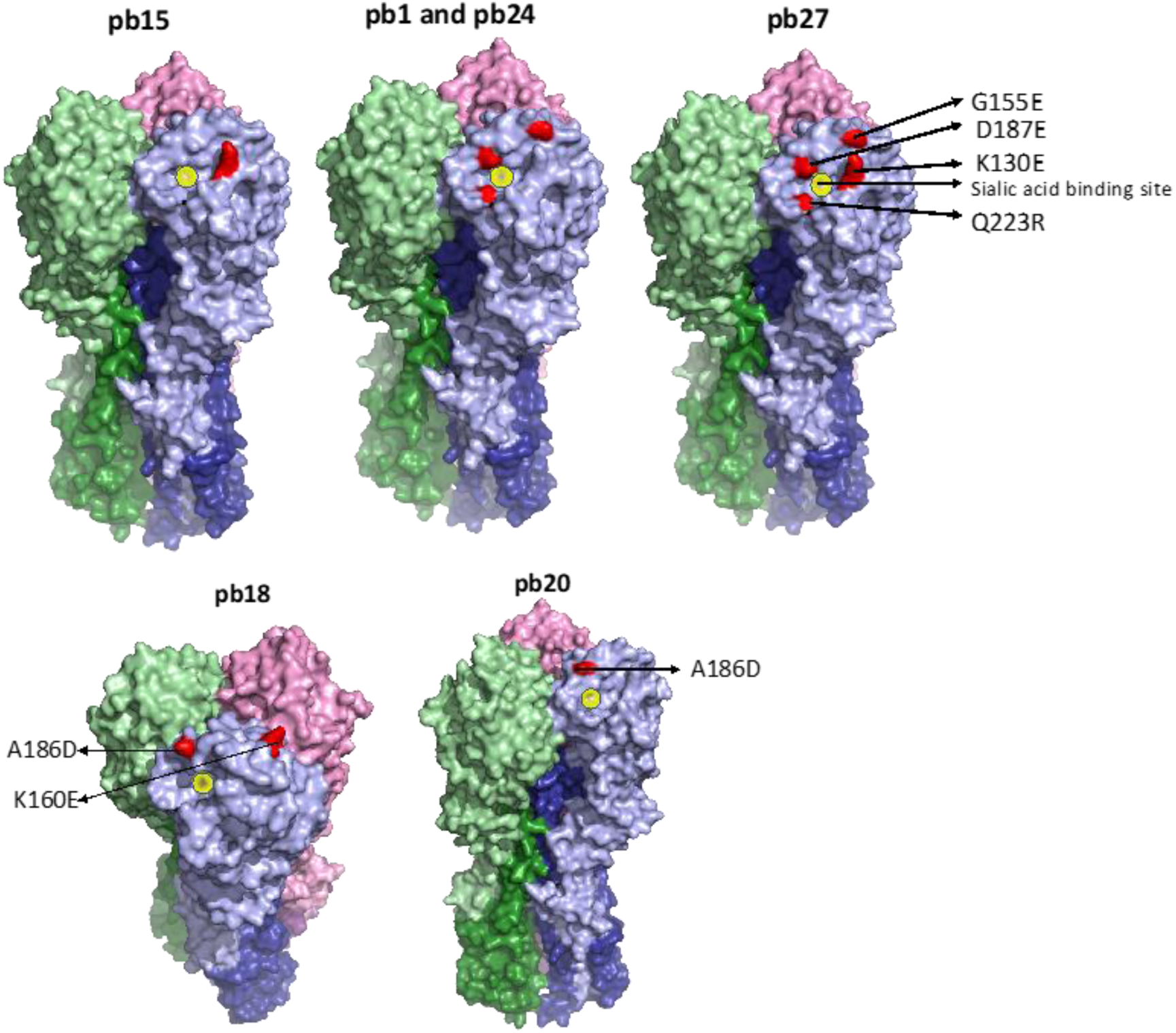
Mapping of the amino acid substitutions selected by mAbs. Substitutions selected by top panel mAbs are - Ca sites (K130E, Q223R) Sa site (G155E) and Sb site (D187E), and on the bottom panel are Sa site (K160E) and Sb site (A186D). Substitutions were mapped with Pymol version 1.7 onto PDB:4M4Y.

Amino acid substitutions at these HA positions have contributed to the antigenic drift of human H1N1pdm09 viruses. Among viruses circulating in 2019-2020, 18% have substitution at K130. While A186 is substituted at a low frequency its neighboring residue, D187, is substituted in 34% of recent viruses. Further, K160M substitution has been found in 3% - 8% of the viruses detected mostly in North America, South America and Australia^27^.

Overall, these results show that nearly a quarter of the porcine mAbs neutralized strongly the H1N1pdm09 virus. The strongly neutralizing mAbs also bind well to MDCK-SIAT1 cells expressing native pHA. In contrast, the remainder of the mAbs bound to recombinant pHA protein in ELISA but showed weak or no binding to the MDCK-pHA cells. Among the high binding and strongly neutralizing mAbs there appear to be two types of porcine mAbs recognizing the two major antigenic sites in pHA: the K163 site (T3-4B or T2-6A like) and the K130 site (2-12C like).

### *In vivo* effect of porcine mAbs

To determine whether porcine mAbs were protective *in vivo*, we administered 10 mg/kg and 1 mg/kg of pb27 intravenously along with a non-treated control group. Unfortunately, two control animals were culled before the start of the experiment for reasons unrelated to the experiment, leaving only three controls. Twenty-four hours later the pigs were challenged with H1N1pdm09 and 4 days later culled to assess virus load and pathology (**Fig. 5a**). The total virus load in nasal swabs was assessed daily over the 4 days. In animals treated with 10 mg/kg the virus load was significantly lower as determined by the area under the curve (AUC) compared to untreated control animals (p= 0.037). At day 1, the difference was also significant (p=0.02) but not on the other days, although there were two pigs out of 5 that did not shed virus at all (**Fig. 5b**). No virus was detected in the BAL and lung in the 10 mg/kg group at 4 DPI. The effect of 10 mg/kg of pb27 is slightly less than the effect of administration of 15 mg/kg of human 2-12C mAb, which we have previously shown in a similar prophylactic experiment^19^. The lower dose (1 mg/kg) of pb27 did not have a statistically significant effect on virus load in nasal swabs or BAL, but no virus was detected in the lung accessory lobe.

**Figure 5.**
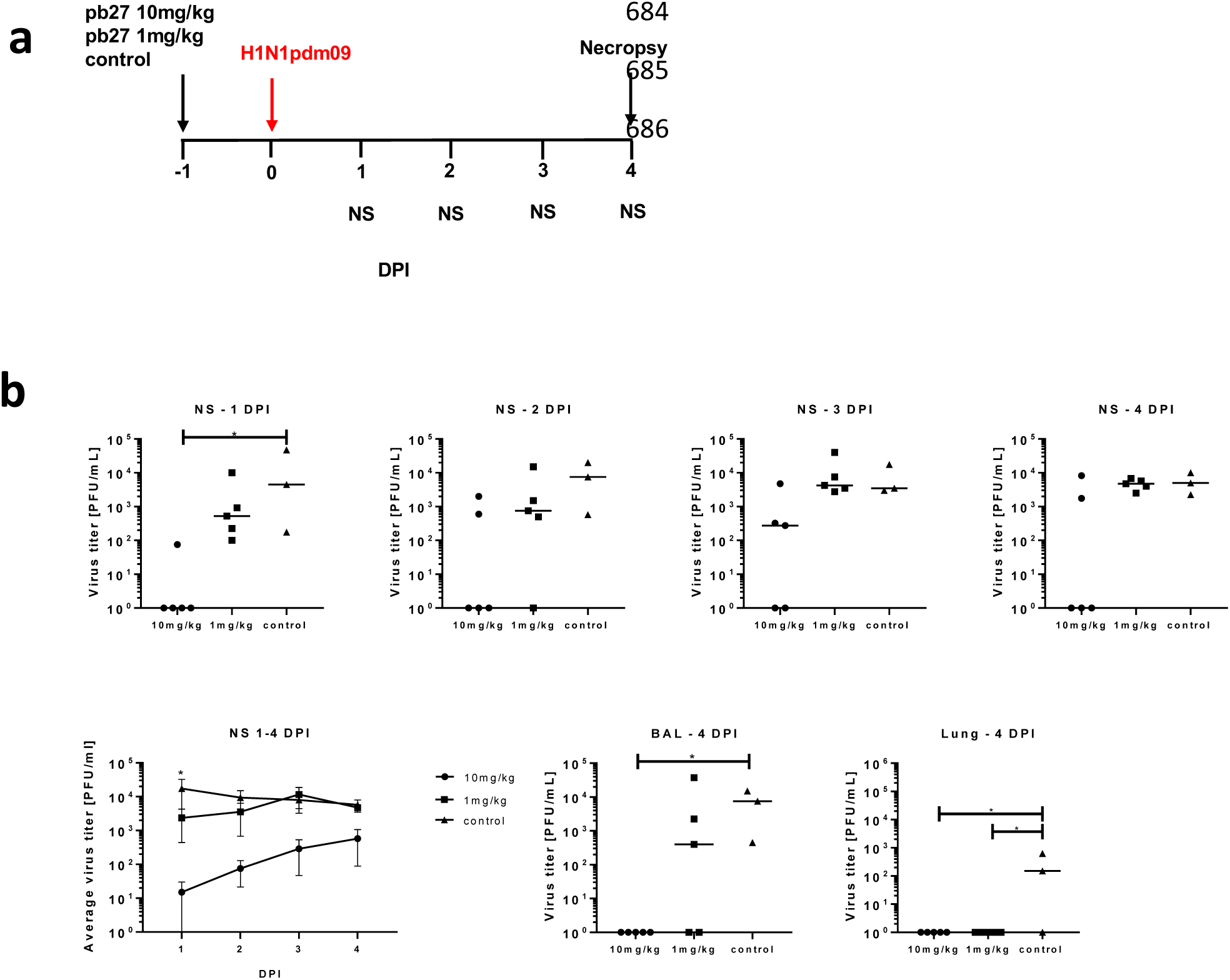
*In vivo* efficacy of pb27. pb27 was administered intravenously to pigs at 10 mg/kg and 1mg/kg, which were infected with H1N1pdm09 virus 24 hours later. Only three control animals were available. Nasal swabs (NS) were taken at 1, 2, 3 and 4 days post-infection (DPI) and pigs euthanized at 4 DPI (**a**). Virus titers in NS, accessory lung lobe (Lung) and BAL at 4DPI were determined by plaque assay (**b**). Each data point represents an individual pig within the indicated group and bars show the mean. Virus shedding in NS is also represented as the mean of the 5 pigs on each day and significance versus diluent control indicated by asterisks. Virus titers were analysed using one-way non-parametric ANOVA, the Kruskal-Wallis test. Asterisks denote significant differences *p<0.05 versus control.

Gross pathology was reduced in both pb27 treated groups, but this was only significant in the 10 mg/kg group (p = 0.003) (**Fig. 6a**). Histopathological evaluation showed typical changes associated with influenza virus infection in the control group, including airway epithelial cell necrosis and attenuation, inflammatory cell infiltration within the airways, alveolar luminae and septae, and perivascular/peribronchiolar lymphoplasmacytic cuffing (**Fig. 6b**). Many bronchiolar and bronchial epithelial cells exhibited positive staining for influenza nucleoprotein (NP). In animals treated with 1 mg/kg, only minimal to mild histopathological changes were observed in 3 of 5 animals, with only a few scattered NP-positive cells within the airway epithelium in 1 of 5 animals from the group. Animals treated with 10 mg/kg showed neither histopathological changes nor NP-stained cells. Two scoring systems used to evaluate the histopathology and compare groups showed significant reduction of histopathology between the 10 mg/kg and the control group (p= 0.0045 for both Morgan and Iowa scores)^28,29^.

**Figure 6.**
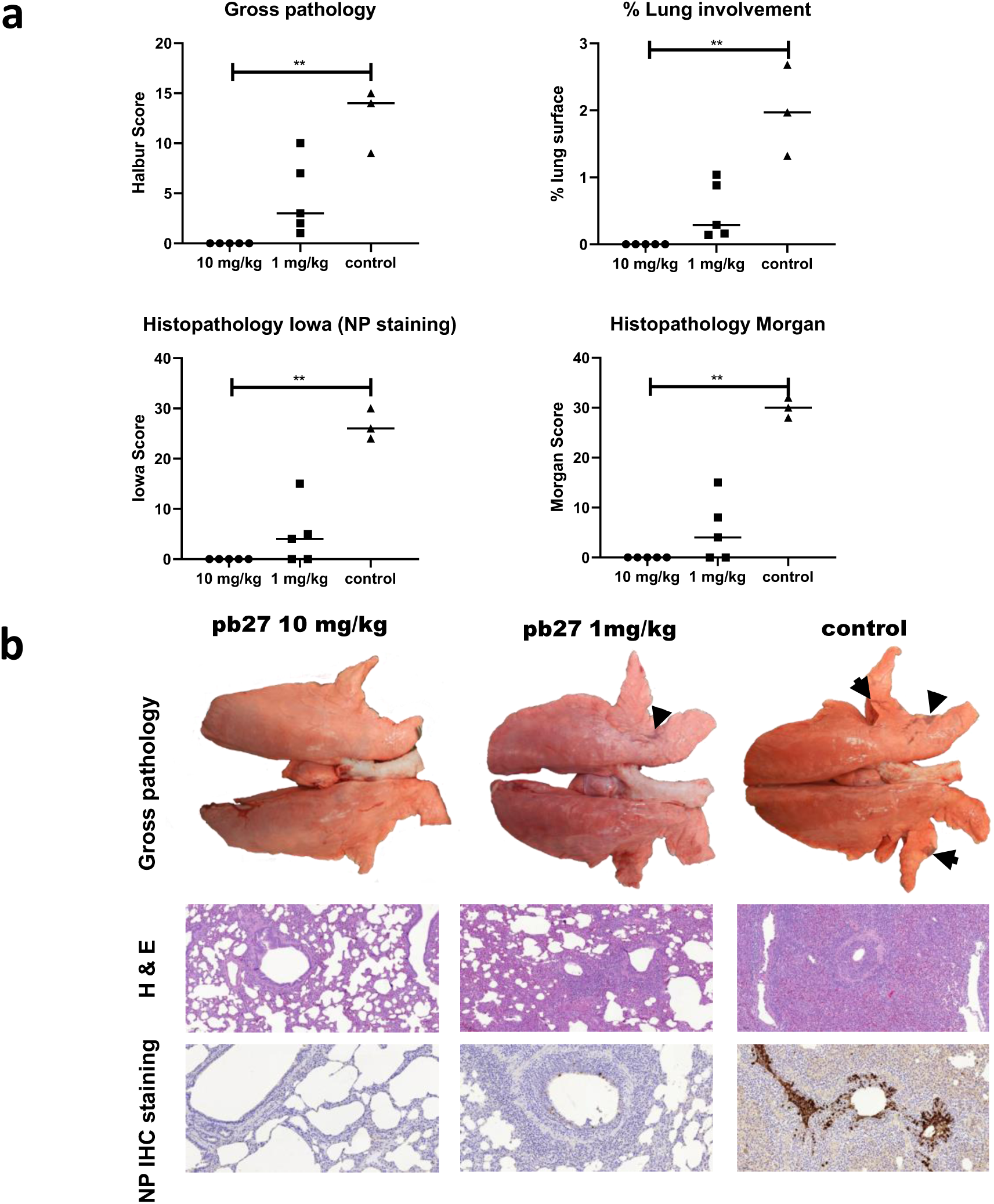
Lung pathology. pb27 was administered intravenously to pigs at 10 mg/kg and 1 mg/kg and animals were infected with H1N1pdm09 virus 24 hours later. The animals were euthanized at 4 DPI and lungs scored for appearance of gross and histopathological lesions. The gross and histopathological scores for each individual in a group and the group means are shown (**a**), including the percentage of lung surface with lesions, the lesion scores and the histopathological scores as previously described (“Iowa” includes the NP staining)^28,29,38^. Representative gross pathology (arrows showing typical areas of focal pneumonia), histopathology (H&E staining; 100x) and immunohistochemical NP staining (200x) for each group are shown (**b**). Pathology scores were analysed using one-way non-parametric ANOVA with the Kruskal-Wallis test. Asterisks denote significant differences *p<0.05, **p<0.01,***p<0.001 versus control.

The concentration of pb27 in porcine serum was determined daily after challenge. A peak concentration of 265 μg/ml pb27 was detected in the 10 mg/kg group and 15 μg/ml in the 1 mg/kg group 24 hours after administration. This declined in both groups over the next 3 days to 65 μg/ml and 7 μg/ml, respectively (**Fig. 7a**). In BAL, pHA-specific antibodies were detected in the pb27 treated groups (mean of 106.5 ng/ml for the 10 mg/kg and 22.4 ng/ml for the 1mg/kg pb27 groups). Neutralizing activity in serum was detected in both 10 mg/kg and 1 mg/kg groups (**Fig. 7b**). There was a 50% inhibition titer of 1:19,200 for the 10 mg/kg group and 1:1,280 for the 1 mg/kg group at the time of challenge. Neutralizing activity in BAL at 1:104 was seen only in the 10 mg/kg group, but the lavage procedure greatly dilutes the fluid present in the airways, although the dilution factor is difficult to calculate.

**Figure 7.**
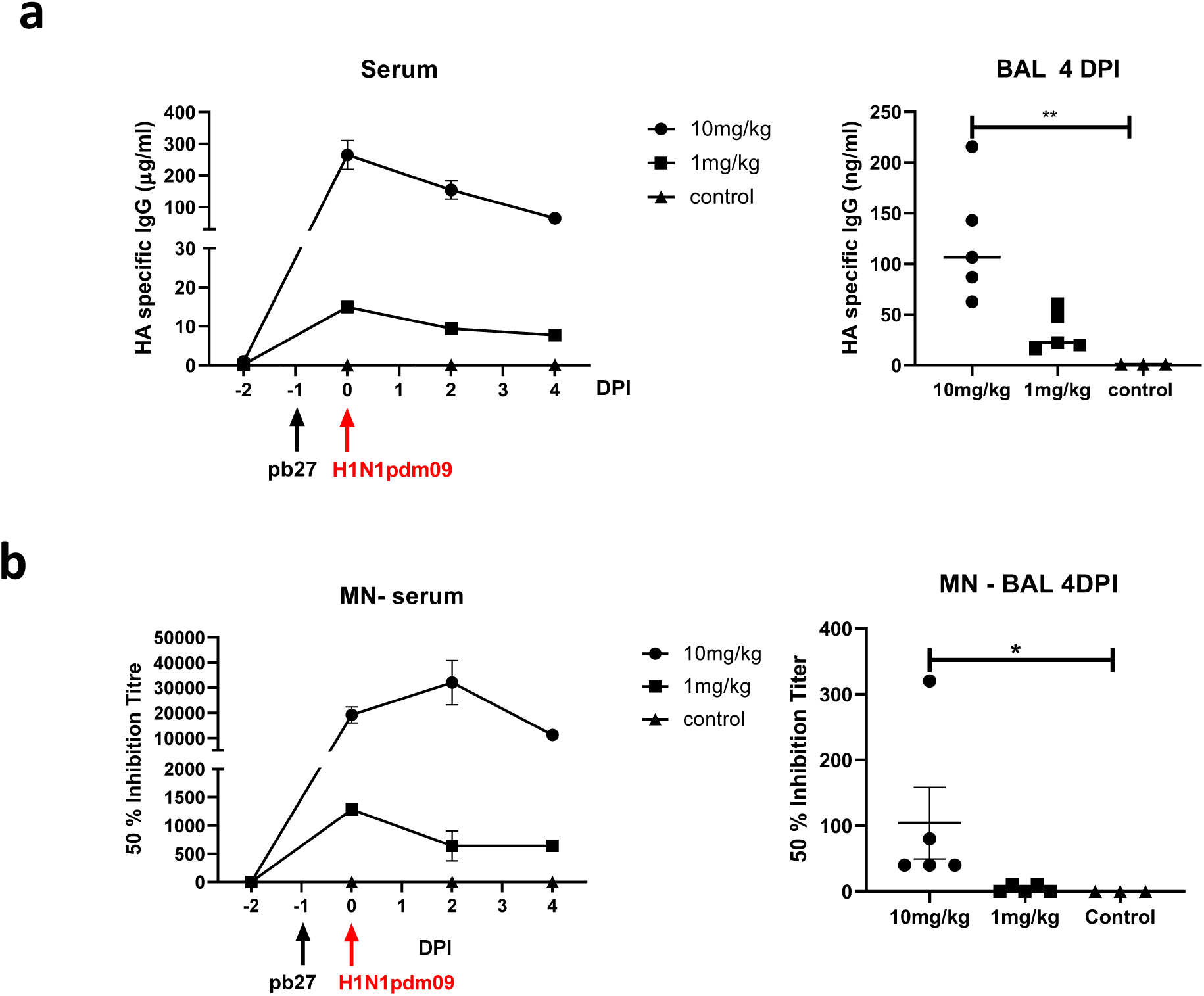
Concentration and neutralizing titer of pb27 in serum and BAL. H1 HA specific IgG in serum, BAL at 4 DPI at the indicated DPI (**a**). 50% neutralization titers against H1N1pdm09 in the serum and BAL at 4 DPI (**b**). Symbols represent individual pigs within the indicated group and lines the mean. Data were analysed using one-way non-parametric ANOVA with the Kruskal-Wallis test. Asterisks denote significant differences *p<0.05 and **p<0.01 versus control.

Overall, these results indicate that pb27 offers robust protection at 10 mg/kg and this correlates with mAb concentration and neutralization activity in serum. Administration of pb27 at 1 mg/kg reduced virus load in the lungs, but not in nasal swabs or BAL. There was a trend towards reduced gross histopathology at 1 mg/kg.

## Discussion

For the first time we have generated porcine influenza specific mAbs from the lung draining lymph nodes of H1N1pdm09 infected pigs. The antibodies are mainly against the HA head and directed towards two major immunodominant epitopes – the Sa site (residues 160 and 163) and Ca site (residue 130), also recognized by humans. The neutralizing activity of these pig mAbs is comparable to the strongest human mAbs. The selected mAb, pb27, reduced virus load and pathology after administration at 10 mg/kg, which resulted in a sustained serum concentration of 100 μg/ml with approximately three logs less in BAL. Lung and BAL virus load was abolished and lung pathology eliminated by 10 mg/kg dosage. Nasal shedding although reduced was not eliminated.

Although pig mAbs against porcine epidemic diarrhoea, classical swine fever and porcine reproductive and respiratory syndrome viruses have been recently generated, here we show that pig mAbs could be produced with high efficiency using draining lymph nodes rather than blood^30-32^. We detected a very low frequency of HA specific B cells in the blood, but a much higher frequency in the local lung tissues and tonsils. Access to draining nodes (by needle biopsy), tonsils and BAL is certainly possible in humans or other large animals and may increase the efficiency of generating mAbs against a variety of disease agents.

The repertoire of IGLV and IGKV gene segments of our mAbs has, as expected for pigs, no apparent usage bias ^21,22^. It is interesting to note that the longest IGH chains tend to pair with IGL; whereas the shortest IGH chains tend to pair with IGK. Also, the IGL chains tend to pair with IGH chains containing a double Cys (so probably a disulphide bond) in the CDRH3. From an evolutionary standpoint, it is interesting to note that cattle almost exclusively use IGL and have longer than usual CDRH3s rich in cysteines^33,34^. However, relatively few clones were analysed in the present study (n = 45) and future work is necessary to determine whether this phenomenon remains true for a larger number of clones across the entire porcine repertoire. We have shown that a strongly neutralizing monoclonal antibody 2-12C against the

HA head administered prophylactically (at 15 mg/kg) to pigs reduced virus load and lung pathology after H1N1pdm09 influenza challenge in a short term experiment^19^. However, longer therapeutic experiments encounter the problem of anti-human antibody responses. Use of pig mAbs will circumvent this difficulty. Although we have not yet measured the pharmacokinetic properties of the porcine mAbs, their availability will allow investigation of how to use mAbs optimally to prevent or treat disease. It is interesting that pb27, a porcine IgG1 mAb, did not abolish virus shedding in nasal swabs. It may be that a different IgG subclass or local administration to the respiratory tract would be more effective in supressing nasal shedding. In our experiment pb27 was given 24h before virus challenge, while in humans mAbs will probably be given post-infection. Further experiments will be needed to determine the efficacy of mAb therapy after infection in the porcine model. The pig influenza model could also be used to evaluate antibody delivery platforms, the effect of IgA and different IgG subclasses and the role of Fc mediated functions. The therapeutic effect of anti-stem and anti-head mAbs and the synergistic effect of cocktail administration could be compared as well.

The specificity of the pig mAbs is very similar to human mAbs known to target H1N1pdm09 HA. They recognised the two major Sa (residues 160 and 163) and Ca (residue 130) sites. In 2014, H1N1pdm09 viruses acquired antigenic drift by substitution at position K163. However, ferret antisera used for monitoring of the viruses failed to detect this antigenic change^27,35,36^. Overall, our results indicate that the pig is an excellent model for understanding how best to apply mAbs as therapy in humans and, since pigs appear to make an antibody response similar to that of humans, pig antisera might be a better surrogate than ferret antisera to detect influenza antigenic drift and thereby inform vaccine recommendations for humans.

## Material and Methods

### Monoclonal antibodies

The pb27 mAb was produced in bulk by Absolute Antibody Ltd (Redcar, UK) and dissolved in 25mM Histidine, 150 mM NaCl, 0.02% Tween, pH6.0 diluent. 1mg each of a further twenty mAbs (pb1, pb5, pb8, pb9, pb11, pb14, pb15, pb16, pb17, pb18, pb20, pb21, pb24, pb27, pb29, pb39, pb31, pb37, pb45, pb46) was also produced by Absolute Antibody Ltd and used in serological assays. The human anti-influenza 2-12C, T2-6A, T3-4B and MEDI8852 mAbs were kindly provided by Alain Townsend.

### Animal studies

The animal experiments were approved by the ethical review processes at the Pirbright Institute (TPI) and the Animal and Plant Health Agency (APHA) and conducted according to the UK Government Animal (Scientific Procedures) Act 1986 under project licence P47CE0FF2. Both Institutes conform to the ARRIVE guidelines.

#### H1N1pdm09 challenge for generation of mAbs

Two 5-to-6 weeks old Landrace x Hampshire cross, female pigs were obtained from a commercial high health status herd and were screened for absence of influenza A infection by matrix gene real time RT-PCR and for antibody-free status by HI using four swine influenza virus antigens - H1N1pdm09, H1N2, H3N2 and avian-like H1N1. Pigs were challenged intra-nasally with 1.5 × 10^7^ PFU of MDCK grown swine A(H1N1)pdm09 isolate, A/swine/England/1353/2009, derived from the 2009 pandemic virus, swine clade 1A.3. (H1N1pdm09) in a total of 4 ml (2 ml per nostril) using a mucosal atomisation device MAD300 (MAD, Wolfe-Tory Medical). Twenty-two days post infection (DPI) the two pigs (pigs 12 and 13) were re-challenged intranasally with 2 × 10^7^ PFU H1N1pdm09. Animals were humanely euthanized 7 days post re-challenge (day 29 post primary challenge) with an overdose of pentobarbital sodium anaesthetic.

#### In vivo efficacy of pb27

Fifteen 5 weeks old Landrace x Hampshire cross, female pigs were obtained from a commercial high health status herd and were screened for absence of influenza A infection and influenza-specific antibody-free status as above. Pigs weighed between 14.5 and 18 kg (average 16.3kg). Pigs were randomized into three groups of five animals as follows: the first group received 10 mg/kg pb27 intravenously via the ear vein after sedation; the second 1 mg/kg pb27 intravenously and the third remained untreated. Two pigs reached their humane end points before the start of the experiment for reasons unrelated to the experiment, leaving only three controls. Twenty-four hours after mAb administration all animals were challenged intranasally with 8 × 10^6^ PFU of H1N1pdm09 in 4 ml (2ml per nostril) using a mucosal atomization device MAD300. Clinical signs (temperature, state of breathing, coughing, nasal discharge, appetite, altered behaviour) observed were mild and none of the pigs developed moderate or severe disease.

### Pathological and histopathological examination of lungs

Animals were humanely euthanized at four DPI with an overdose of pentobarbital sodium anaesthetic. The gross and histopathological analyses was performed as previously described^29,37^. Briefly, lungs were removed and digital images were taken of the dorsal and ventral aspects. Gross pathology was scored as previously described^38^ and the lung surface with lesions was calculated by digital image analysis using Nikon NIS-Ar software. Lung tissue samples were taken from the apical, medial and diaphragmatic lobes and fixed in 10% buffered formalin. Four micron sections were cut and stained with H&E for histopathological analysis and with IHC using a mouse mAb against the virus nucleoprotein (NP)^37^. Histopathological lesions were scored by a qualified veterinary pathologist following two systems previously described^28,29^.

### Tissue Sample Processing

Two nasal swabs (one per nostril) were taken daily following infection with H1N1pdm09. Tonsil, mesenteric, tracheobronchial (TBLN) and mandibular lymph nodes were dissected out, cut into smaller pieces and tissue integrity disrupted by squashing them with the back of a plunger in the presence of RPMI supplemented with 10% FBS. The tissue homogenates were passed through 100 µM cell strainers (Sigma, UK). Blood, spleen, broncho-alveolar lavage (BAL) and accessory lung lobe were processed as described previously ^29,37^. Virus titer in nasal swabs, BAL and lung was determined by plaque assay on MDCK cells as previously described^29^.

### Single cell sorting

Cryopreserved single cell suspensions from the TBLN were thawed and rested for 1 to 2h at room temperature before staining for surface markers and HA binding. For the single cell sort 1.4 × 10^7^ cells from pig 12 were stained with CD3-RPE (Clone: PPT3, BIO-RAD, UK), CD8α-RPE (Clone: MIL12, BIO-RAD, UK), CD172α-RPE (Clone: 74-22-15, BIO-RAD, UK) and 50 µg/ml biotinylated, trimeric H1N1pdm09 (A/England/195/2009) HA (pHA) for 30 min at 4°C in the dark. After the incubation step, cells were washed with PBS twice before staining with the following antibodies or reagents for another 30 min at 4°C in the dark: Streptavidin-BV650 (Biolegend, UK), IgG (H+L) AF 647 (Mouse anti-pig IgG (H+L) AF 647, Generon, UK) and near-IR fixable Live/Dead stain (Invitrogen, UK). After the final labelling steps, the cells were washed 3 times in PBS and then re-suspended in 0.5 ml of pre-chilled 0.5% BSA PBS and passed through a 70 μm cell strainer (BD Biosciences, UK) prior to cell sorting. Single colour controls were used for compensation and fluorescence minus one primary Ab controls were used to set thresholds.

pHA-specific B cells were identified and collected, using DIVA 8 acquisition software and a FACS Aria III cell sorter (BD Biosciences), at single cell density into hard shell 96-well PCR plates (BIORAD, UK) containing 10 μl/well of RNA catch buffer (10mM Tris pH7.4 supplemented with RNasin (Promega, UK)). Full 96 well plates were immediately covered with aluminium foil seals, centrifuged for 5 min at 300g, 4°C and transferred to -80°C. The FACS gating strategy to identify HA positive, single B cells is shown in **Suppl Fig 1**. In brief, samples were gated on lymphocytes (SSC-A vs FSC-A) and singlet (FSC-H vs FSC-A) live cells were identified by a live/dead stain. HA-specific IgG^HI^ cells were then determined as CD3-, CD8α-, CD172α- and double positive for HA and IgG.

### Cloning of light and heavy chains from single cells

We used a two-step RT-PCR method to amplify the variable genes of the heavy and light chains. cDNA was synthesized in one reaction and aliquots of the cDNA were used in subsequent PCRs to amplify VDJ heavy or VJ light genes in separate reactions. cDNA synthesis on single, sorted cells was carried out in a total reaction volume of 20 µl using Sensiscript reverse transcriptase (Qiagen, UK) containing a mix of oligonucleotides dT_23_VN (NEB, UK) and random hexamer primers (NEB, UK). The cDNA reaction was incubated for 60 min at 40°C. Two to 4µl of cDNA was used to amplify the heavy and light chains respectively. For amplification of the heavy and kappa chains a nested PCR approach was chosen to have enough material for the downstream cloning whereas for the lambda chain one PCR yielded enough of the PCR product for the subsequent cloning step. For all PCRs, the Q5 High-Fidelity 2x Master Mix (NEB, UK) was used according to the manufacturer’s instructions in a total reaction volume of 40 µl. The following oligonucleotide primers were used for the first PCR of the heavy chains IGHV_L1_F, IGHV_L2_F and IGHG_191R (see supplementary table 1) which amplify all IgG subtypes. The products of the PCR were analysed on a 1% agarose/1xTAE gel. If there was only one specific product, the PCR product was purified with the GFX PCR DNA and Gel Band Purification Kit (GE Healthcare, UK) and eluted in dH_2_O. In case of multiple products, the right sized product was extracted, purified and eluted in dH_2_O.

Up to 20 ng of DNA was used as template for the second PCR with subtype-specific reverse primers (IgG1_R_HiFi or IgG3_R_HiFi) and the IgG forward oligo, IgG_F_HiFi that binds in the conserved FR1 of the V gene of IGH (**Suppl Table 1**).

The second PCR product (V, D, J heavy genes) was cloned in frame by HiFi assembly (NEB, UK) into the *Kpn*I/*Pst*I (NEB, UK)-linearized expression vector containing the IgG1 constant domain. The kappa VJ region was also amplified with a nested PCR approach using IgK_L1_F, IgK_L2_F and IgK_R primers in the first PCR reaction. The PCR products were either gel extracted or directly purified and used as the template for the second PCR with primers IgK_V1_F_HiFi IgK_V2_F_HiFi and IgK_R_HiFi. The products from the second PCR were cloned into the kappa expression vector.

Amplification of the lambda chain was successfully achieved in one 40 µl PCR reaction using IgL_V3_F_HiFi, IgL_V8_F_HiFi and IgL_R_HiFi primers (**Suppl Table 1**). The purified product was used directly for the HiFi assembly into the porcine lambda expression vector. The expression vectors pNeoSec-SsFc-IgG1, pNeoSec-SsLC-κ and pNeoSec-SsLC-λ were provided by the Immunological Toolbox (The Pirbright Institute, UK). The vectors are derived from pNeoSec^39^. They encode the µ-phosphatase secretion leader and were modified to include the antibody v region coding sequence cloning site containing the lacZ promoter and lacZ alpha peptide flanked by *KpnI* (5’) and *PstI* (3’), and the porcine constant domains of IgG1, kappa or lambda), as previously described for expression of recombinant mouse antibodies ^40^. The heavy and light chain PCR products were cloned in frame with seamless cloning using the NEBuilder® HiFi DNA Assembly Master Mix (NEB, UK) after linearizing the vectors with *KpnI* (5’) and *PstI* (3’). The forward oligos had a 5’-20bp-overlap specific for the leader sequence encoded in the expression vector for HiFi assembly, whereas the reverse oligos all primed in the 5’ end of the constant domains thereby also generating an overlap with the expression vectors suitable for seamless cloning.

The heavy and light chain assembly reactions from each single cell were transformed into NEB® 5-alpha Competent E. coli (High Efficiency) (NEB, UK). Between 2 to 6 colonies from each transformation were picked and inoculated in 3 ml LB + Kanamycin overnight aerobically at 37°C on an orbital shaker. Plasmid DNA was isolated with the QIAprep Spin Miniprep Kit (Qiagen, UK) and submitted to Sanger sequencing in-house (The Pirbright Institute, UK) to ensure an intact and full-length open reading frame before transfecting into HEK293 cells.

### Expression and purification of porcine antibodies

HEK 293 cells (provided by Cell Culture Unit (CCU), The Pirbright Institute, UK) were seeded in T25 cm^2^ tissue culture flasks in RPMI supplemented with HEPES, Glutamax and 10% FBS the day before transfection. Four µg of the heavy chain and 4 µg of the corresponding light chain were mixed together in 375 µl Opti-MEM (Gibco, ThermoFisher Scientific, UK) before addition of polyethylenimine (PEI MAX (Polysciences, Generon, UK) in a 3:1 ratio (PEI:DNA) in an equal volume of OptiMEM. Before transfection the medium was changed to RPMI supplemented with HEPES, Glutamax and 2% FBS. After 20 min the DNA:PEI complex was added dropwise onto the HEK 293 cells. After 4h the transfection mix-containing medium was discarded, cells were washed once with sterile PBS and cultured for up to 4 days in RPMI supplemented with HEPES, Glutamax, Pen/Strep and 5% IgG-depleted FBS (CCU, The Pirbright Institute or GIBCO, ultra-low IgG, ThermoFisher Scientific, UK).

After 5 days culture supernatant was harvested and cleared of cells and debris by centrifugation for 10 min at 500 x g. Thirty µl of Protein G (Protein G Sepharose Fast Flow, Merck, UK) slurry per sample was added in addition to 0.01% sodium azide and incubated on a roller overnight at 4°C. After incubation, samples were centrifuged at low speed, the supernatant was discarded and the Protein G beads were resuspended in sterile PBS and transferred into Spin columns (Micro Bio-Spin Chromatography Columns, Bio-Rad, UK). The beads were washed 4 x with sterile PBS before addition of 117 µl Glycine pH 2.7 per sample and incubated for 5 min at room temperature. After that mAbs were eluted into sterile microcentrifuge tubes prefilled with 13 µl of 1M Tris pH 8.0 and the buffer was exchanged to PBS/0.01% NaN_3_ using Zeba Spin Desalting Columns (ThermoFisher Scientific, UK) according to the manufacturer’s instructions and stored at 4°C until further analysis.

### B cell ELISpot

B cell ELISpots were performed for the detection and enumeration of antibody-secreting cells in single cell suspensions prepared from different tissues and peripheral blood collected from two pigs 7 days after re-challenge with H1N1pdm09. ELISpot plates (Multi Screen-HA, Millipore, UK) were coated with 100 μl per well of appropriate antigen or antibody diluted in carbonate/bicarbonate buffer for 2h at 37°C. To detect HA-specific spot-forming cells, plates were coated with 2.5 µg per well of recombinant pHA from H1N1pdm09 (A/England/195/2009) for the enumeration of total IgG-secreting cells with 1 µg per well of anti-porcine IgG (mAb, MT421, Mabtech AB, Sweden) or with culture medium supplemented with 10% FBS (background). The coated plates were washed with PBS and blocked with 200 µl/well 4% milk in PBS. Freshly isolated or frozen porcine cell suspensions from different tissues were filtered through sterile 70 µM cell strainers, plated at different cell densities in culture medium (RPMI, 10% FBS, HEPES, Sodium pyruvate, Glutamax and Penicillin/Streptomycin) on the ELISPOT plates and incubated for a minimum of 18 h at 37°C in a 5% CO_2_ incubator. After incubation the cell suspension was removed, the plates washed once in ice-cold sterile H_2_O and thereafter with PBS/0.05 % Tween 20, before incubation with 100 µl per well of 0.5 µg/ml biotinylated anti porcine IgG (mAb, MT424, Mabtech AB, Sweden) diluted in PBS/0.5 % FBS for two hours at room temperature. Plates were washed with PBS/0.05% Tween 20 and incubated with streptavidin – alkaline phosphatase conjugate (Strep-ALP, Mabtech AB, Sweden). After a final wash, the plates were incubated with AP Conjugate Substrate (Bio-Rad, UK) for a maximum of 30 min. The reaction was stopped by rinsing the plates in tap water and dried, before spots were counted.

### Serological assays

ELISA was performed using the recombinant pHA containing a C-terminal thrombin cleavage site, a trimerization sequence, a hexahistidine tag and a BirA recognition sequence as previously described^27^. Hemagglutination inhibition (HAI) Ab titers were determined using 0.5% chicken red blood cells and H1N1pdm09 at a concentration of 4 HA units/ml. Microneutralization (MN) was performed using standard procedures as described previously^19,41^.

The porcine mAbs were also tested for binding to MDCK-SIAT1 cells stably expressing pHA from H1N1pdm09 (A/England/195/2009), HA from A/Puerto Rico/8/1934 (PR8, H1N1) and H5 HA (A/Vietnam/1203/2004). Confluent cell monolayers in 96-well microtiter plates were washed with PBS and 50 ul of the antibody dilution in PBS/0.1% BSA was added for 1 h at room temperature. The plates were washed three times with PBS and 100 ul of horseradish peroxidase (HRP)-conjugated goat anti-pig Fc fragment secondary antibody (Bethyl Laboratories diluted in PBS, 0.1% BSA) was added for 1 h at room temperature. The plates were washed three times with PBS and developed with 100 µl/well TMB High Sensitivity substrate solution (Biolegend). After 5 to 10 min the reaction was stopped with 100 µl 1M sulfuric acid and the plates were read at 450 and 570 nm with the Cytation3 Imaging Reader (Biotek). The cut off value was defined as the average of all blank wells plus three times the standard deviation of the blank wells.

### Selection of influenza variants with monoclonal antibodies

Approximately 0.5 × 10^7^ TCID_50_ X-179A virus was mixed with antibody at 10 µg/ml in a total volume of 1 ml topped with virus growth medium (DMEM, 0.1%BSA, 10 mM HEPES, pH7.0, penicillin and streptomycin) and incubated at 37°C for 1 hour. The virus and antibody mixture were added to a monolayer of MDCK-SIAT1 cells in a 6-well plate. After 40 min, 2 ml of virus growth medium with 1.5 µg/ml TPCK trypsin was added to the cells and incubated for 2 days at 37 °C with 5% CO_2_. Virus in the culture supernatant was harvested and tested in MDCK-SIAT1 cell infection assays. The HA and NA genes of the escape mutant virus populations were sequenced to assess the presence of encoded amino acid substitutions compared to the X-179A parent virus.

### Competitive Binding Assay

For epitope mapping, the inhibition of binding of known human mAbs 2-12C (head), T3-4B (head) and MEDI8852 (stem) at 1 µg/ml was performed with 10 µg/ml porcine mAbs. The blocking of human mAb binding to pHA expressed on the surface of MDCK-SIAT1 cells was detected using second layer goat-anti-human IgG Alexa fluor 647 (Invitrogen A-21445) for 2-12C or streptavidin-Alexa fluor 647 (Invitrogen S21374) for biotin-labelled T3-4B and MEDI8852. Fluorescence was measured using Clariostar (BMG Labtech) and the binding (%) of human mAb was calculated as (X-Min)/(Max-Min)*100 where X = Measurement of the competing mAb, Min = PBS background, Max = Binding of huMab in presence of non-binding mAb. Means and 95% confidence intervals for eight replicates are shown.

## Supplementary information

**Supplementary Fig. 1**: Gating strategy for isolating single HA specific antibody secreting cells.

**Supplementary Table 1:** Primer sequences

## Acknowledgements

We are grateful to the animal staff for providing excellent animal care and to the immunological toolbox for providing the porcine heavy and light chain Ig constructs. We thank Giacomo Gorini, Carolyn Nielsen and Simon Draper from the Jenner Institute, University of Oxford for helpful discussions. We thanks APHA for providing the swine A/Sw/Eng/1353/09 influenza virus strain (DEFRA surveillance programme SW3401).

## Funding

This work was supported by Bill & Melinda Gates Foundation grant OPP1201470 and OPP1215550 (Pirbright Livestock Antibody Hub); UKRI Biotechnology and Biological Sciences Research Council (BBSRC) grants BBS/E/I/00007031, BBS/E/I/00007038 and BBS/E/I/00007039. P.R. and A.R.T. were funded by the Chinese Academy of Medical Sciences (CAMS) Innovation Fund for Medical Sciences (CIFMS), China Grant 2018-I2M-2-002, the Townsend-Jeantet Prize Charitable Trust (charity number 1011770) and the Medical Research Council (MRC) Grant MR/P021336/1.

## Author contributions

ET, BH, AM, PR, ART, JH conceived, designed and coordinated the study. ET, AM, BH, BP, BC, MP, EB, TM, KM, MBDP, SPG, PB, RMR, JCS, ART, WM designed and performed experiments, processed samples and analyzed the data. JWM and RSD performed sequencing analysis. FJS carried out postmortem and pathological analysis. ET, PR, AM, BH, PB, ART wrote the manuscript. All authors read and commented on the manuscript.

## Competing interests

The authors have no financial conflicts of interest.

## Notes

### Competing Interest Statement

The authors have declared no competing interest.

