## Supplemental table and figure for "Protective porcine influenza virus-specific monoclonal antibodies recognize similar haemagglutinin epitopes as humans"

### Supplementary Table 1: Primer sequences

| Primer | Sequence (5' to 3') | Use | Target sequence |
|---|---|---|---|
| IGHV_L1_F | AACTGGGTGGTCTTGTTTGC | 1st PCR IGH | Leader sequence IgG |
| IGHV_L2_F | TCTCTTACAAGGTRTCCAGGGTG | 1st PCR IGH | Leader sequence IgG |
| IGHG_191R | GGAGTAGAGCCCTGACGG | 1st PCR IGH | Conserved region in constant domain of all IgG isotypes |
| IgG1_R_HiFi | CGATGGGGCCGTCTTGG | 2nd PCR HiFi assembly | IgG1 constant domain |
| IgG_R_HiFi | tgctgatgggttgcgtagctGAGGAGAAGCTGGTGGAGTCTG | 2nd PCR HiFi assembly | FR1_IgG |
| IgG3_R_HiFi | GTAGACCGATGGAGCTGTGTTGT | 2nd PCR HiFi assembly | IgG3 constant domain |
| IgL_V3_F_HiFi | tgctgatgggttgcgtagctTATGAGCTGACCCAGCCGTC | PCR HiFi assembly (Lamda) | Binds in FR1 of IgL_V3 |
| IgL_V8_F_HiFi | tgctgatgggttgcgtagctCAGACTGTGATCCAGGAGCC | PCR HiFi assembly (Lamda) | Binds in FR1 of IgL_V8 |
| IgL_R_HiFi | GGAGCGGCCTTGGGCT | PCR HiFi assembly (Lamda) | Binds in constant region of IgL |
| IgK_L1_F | GCCTCYTGCTGCTCTGG | 1st PCR (Kappa) | Binds in leader of IgK |
| IgK_L2_F | TTCCCTGCTCAGCTCCTG | 1st PCR (Kappa) | Binds in leader of IgK |
| IgK_R | CTAAGCCTCACACTCGTTCCTG | 1st PCR (Kappa) | Binds in constant region of IgK, 3'end |
| IgK_V1_F_HiFi | tgctgatgggttgcgtagctGCCATCCAGCTGACCCAG | 2nd PCR HiFi assembly (Kappa) | Binds in FR1 |
| IgK_V2_F_HiFi | tgctgatgggttgcgtagctGCCATYGTGCTGACCCAG | 2nd PCR HiFi assembly (Kappa) | Binds in FR1 |
| IgK_R_HiFi | ACGGATGGCTTGGCATCAGC G | 2nd PCR HiFi assembly (Kappa) | Binds in constant region of IgK, 5' end |

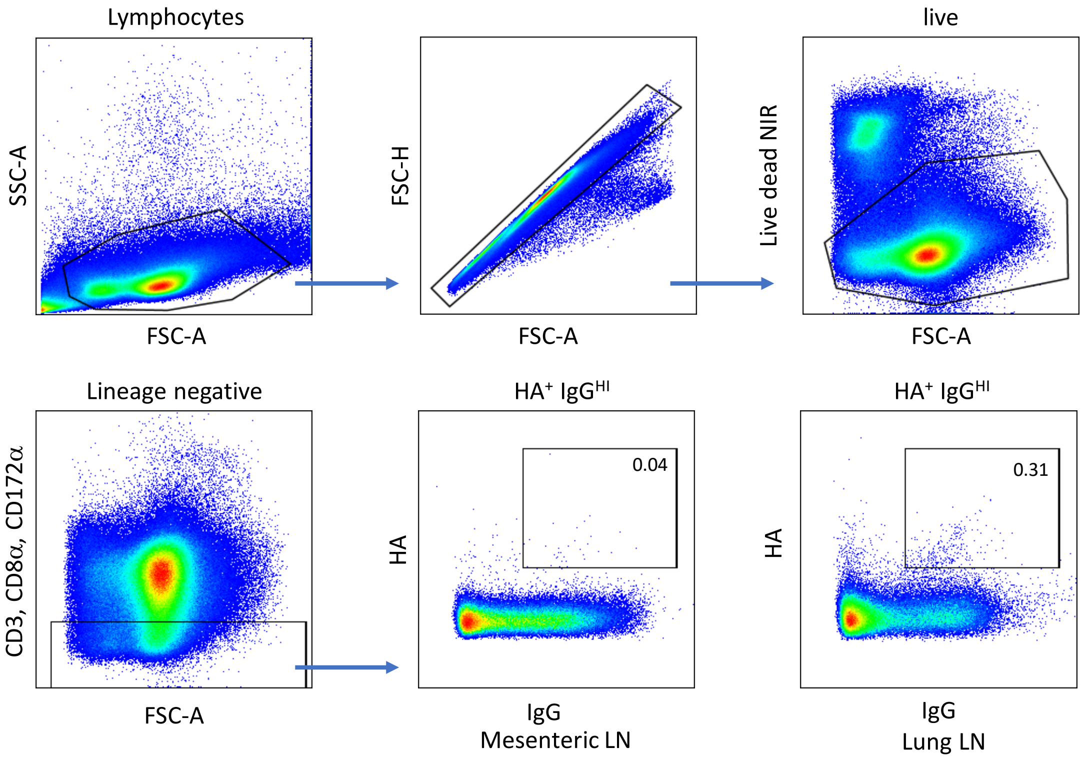


**Supplementary Figure 1: Gating strategy for isolating single HA specific antibody producing cells.** Samples were gated on lymphocytes (SSC-A vs. FSC-A) and singlets (FSC-H vs. FSC-A), live cells were identified as negative for live dead stain. HA-specific $IgG^{HI}$ cells were then identified as CD3-, CD8α-, CD172α- and double positive for HA and IgG.
